# Hydroxyl and Trifluoromethyl Radical Carbohydrate Footprinting for Probing Protein Binding Components of Oligosaccharides

**DOI:** 10.1101/2025.05.06.652515

**Authors:** Quadrat Yusuph, Sandeep K. Misra, Hao Liu, Joshua S. Sharp

## Abstract

Carbohydrates are found in various forms in living organisms, both as free-standing glycans as well as glycoconjugates including glycoproteins, glycolipids, and glycosaminoglycans. These structures play crucial roles in many biological processes, often mediated or influenced by interactions of carbohydrates with other biomolecules. However, studying these interactions is particularly challenging due to the structural complexity of carbohydrates, their dynamic conformational behavior, and the low binding affinities often involved. To address these challenges, we are developing a novel method that leverages mass spectrometry-based radical footprinting of carbohydrates (RFC). We monitored changes in the solvent accessibility of specific regions within oligosaccharides by measuring variations in the apparent rate of hydroxyl radical and trifluoromethyl radical-mediated oxidation. In our studies, a collection of trisaccharide isomers and N, N′,N′′-triacetylchitotriose (NAG_3_) shows no significant change in modification in non-binding protein solutions. However, in the presence of two proteins that bind NAG_3_ specifically, NAG_3_ oxidation is reduced. We find that the free reducing end is the primary site of hydroxyl radical oxidation under covalent labeling conditions, allowing it to distinguish interactions at the glycan reducing end. Trifluoromethyl radicals, conversely, label broadly across the trisaccharide by substitution into a C-H bond. Overall, this approach offers a powerful new approach for identifying glycan-protein interactions and mapping the binding interface of glycans.

## INTRODUCTION

Carbohydrates play diverse roles in biological systems. They serve as primary energy source and form structural components like cellulose in plant cell walls and chitin in fungi and arthropod exoskeletons. Glycoconjugates are broadly found on the cell surface, playing a variety of roles in cellular biology. Structurally, carbohydrates range from simple monosaccharides like glucose to complex polysaccharides, with complexity arising from anomeric configurations, linkage positions, branching, and modifications [1, 2]. There are many types of carbohydrates on the mammalian cell surface, including *N*-glycans, *O*-glycans, glycosaminoglycans (GAGs), lipo-oligosaccharides and glycolipids. These carbohydrates bind with various proteins and mediate various biological processes [3]. These interactions can be monovalent or multivalent and are often dynamic. Understanding carbohydrate interactions is essential to understanding biology across various scientific domains due to their roles in cellular communication, pathogen-host interactions, disease development, drug delivery, biotechnology, nutritional science, and structural biology.

Studying carbohydrate interactions from complex mixtures is fraught with challenges due to the structural complexity, diversity, and dynamic nature of carbohydrates. Glycan microarrays offer high-throughput screening but face hurdles in representing the full natural complexity of glycan structures. Various biophysical and computational methods have been utilized to study carbohydrate interactions from different perspectives. Techniques such as X-ray crystallography [4], NMR spectroscopy [5, 6], molecular modeling [7], surface plasmon resonance [8], and isothermal titration calorimetry [9, 10] have all contributed to uncovering critical structural details involved in the formation of carbohydrate-protein complexes. They are limited by the amount of sample needed and the purity of the sample required.

MS is now commonly used to study glycan-protein interactions. Electrospray ionization (ESI) MS directly measures the free and glycan-bound protein ions in solution [11]. One promising approach screens interaction based on a catch-and-release-ESI-MS assay, where the binding glycan first “catches” the protein and then is “released” by CID and identified by MS [12]. A filter-entrapment enrichment pull-down assay coupled with MS analysis has been used to study GAG-protein interactions in a rapid screening manner, including various binding affinities [13]. Overall, MS-based characterization approaches are extremely useful. However, current MS-based technologies generally cannot probe the specific portions of the glycan involved in the interaction. Protein-carbohydrate interactions have been extensively studied using footprinting techniques, but only from the perspective of the protein, identifying protein surfaces that interact with the carbohydrate [14-18]. There is currently no established footprinting methodology for carbohydrates. In 2014, a novel approach was introduced for fingerprinting polysaccharides through reaction with hydroxyl radicals (•OH). Their work demonstrated that hydroxyl radicals can induce non-enzymatic scission in polysaccharides, particularly pectin, using a fluorescent labeling technique to document oxidative damage [19]. Given the diverse composition of glycan preparations and the vital role of mass spectrometry in glycan research, glycan footprinting would be a valuable tool for investigating glycan interactions with other biomolecules.

Here, we report the footprinting of a model trisaccharide, triacetylchitotriose (NAG_3_), by both hydroxyl and trifluoromethyl radicals. Hydroxyl radical oxidation of NAG_3_ was performed using excimer laser photoexcitation of hydrogen peroxide [20] and footprinting by trifluoromethyl radicals was performed by hydroxyl radical-induced decomposition of sodium triflinate [21] using the Flash Oxidation (FOX) Photolysis System, a commercially available standardized system that utilizes a broadband UV flash lamp [22].

## MATERIALS AND METHODS

### Materials

Triacetylchitotriose (NAG_3_), 1-kestose, isomaltotriose, raffinose, melezitose, myoglobin, ubiquitin, tris base, dimethylthiourea, sodium triflinate, and sodium borohydride were bought from Sigma (St. Louis, MO). Disodium phosphate and monosodium phosphate were purchased from VWR (Radnor, PA). LC-MS-grade water, acetonitrile, formic acid, and hydrogen peroxide were purchased from Fischer Chemicals (Fair Lawn, NJ). Lysozyme was bought from Goldbio (St. Louis, MO). Griffonia lectin was purchased from Vector Laboratories Inc. (Burlingame, CA). Methionine amide was obtained from Bachem (Torrance, CA). Sequencing-grade-modified trypsin was purchased from Promega Corp. (Madison, WI). We used all reagents without further purification. All pH measurements were made with a Mettler Toledo pH meter at 25^°^C.

### Hydroxyl radical carbohydrate footprinting

A glycan mixture containing 25 μM each of NAG_3_, isomaltotriose, 1-kestose, raffinose, and melezitose was prepared in 10 mM sodium phosphate buffer (pH 7.4). Four proteins were evaluated in this study, comprising two non-carbohydrate-binding proteins (ubiquitin and myoglobin) and two carbohydrate-binding proteins (lysozyme and lectin). Each protein was prepared in combination with the glycan mixture at a glycan-to-protein ratio of 1:2, with the proteins also dissolved in 10 mM sodium phosphate buffer (pH 7.4) at a final concentration of 50 μM. The hydroxyl radical carbohydrate footprinting (HRCF) experiment was performed based on established protocols [23, 24]. Briefly, a mixture containing 1 mM adenine, 17 mM glutamine, and 100 mM hydrogen peroxide was loaded into a gastight syringe (Hamilton, Reno, NV). The mixture was passed through the focused beam path of a COMPex Pro 102 KrF excimer laser (Coherent, Inc., CA) operating at 20 Hz with an exclusion volume of 15% and a fluence of approximately 13 mJ/mm^2^. 20 μL of exposed samples were collected into a microcentrifuge tube of 25 μL of quenching solution containing 0.3 μg/μL catalase and 0.5 μg/mL methionine amide. Adenine absorbance at 260 nm was measured using a NanoDrop UV/VIS spectrophotometer (Thermo Scientific) to confirm consistent radical dose delivery across all experimental replicates. This measurement ensured the reproducibility of the effective radical exposure. All the samples were prepared and analyzed in triplicate.

### Trifluoromethyl radical carbohydrate footprinting

The reaction conditions were optimized using NAG_3_ as the model compound by varying the concentration of hydrogen peroxide across a range of 10 to 100 mM at a fixed concentration of 40 mM sodium triflinate. NAG_3_ was dissolved in LC-MS water to achieve a final concentration of 25 μM. We generated trifluoromethyl radicals in situ using 40 mM of sodium triflinate, CF_3_SO_2_Na (Langlois Reagent-LR), and 10 mM hydrogen peroxide. Hydrogen peroxide was added immediately before injection, and samples were illuminated with a Fox Photolysis system [25] at 900 V lamp voltage (GenNext Technologies, Inc., Half Moon Bay, CA). A combination of quench and reduction solution containing 1 M sodium borohydride, 70 mM methionine amide, and 100 mM dimethylthiourea (DMTU) of an equal volume as the illuminated sample was placed in the collection tube prior to illumination. We deposited the sample into this quench/reduction solution immediately after photolabeling. After illumination and quenching, we incubated the samples at 37 ^°^C with rotation for 20 hours.

We used five different glycans (25 μM each of NAG_3_, isomaltotriose, 1-kestose, melezitose, and raffinose) and four different model proteins (lysozyme, lectin, myoglobin, or ubiquitin). Each protein was prepared in combination with all five glycans at a glycan-to-protein ratio of 1:2, with each glycan-protein combination prepared in a separate microcentrifuge tube. Each protein was dissolved in 10 mM sodium phosphate buffer (pH 7.4) to achieve a final protein concentration of 50 μM. We generated trifluoromethyl radicals in situ using 40 mM of sodium triflinate (CF3SO2Na, Langlois Reagent) and 40 mM of hydrogen peroxide. The illumination and quenching procedures were performed as described in the previous section, and the samples were stored at -20 °C until analysis by liquid chromatography-mass spectrometry (LC-MS).

### HDX-Direct Infusion MS of NAG_3_

NAG_3_ was dissolved in LC-MS grade water to a final concentration of 25 μM. Trifluoromethyl radicals were generated in situ using 40 mM sodium triflinate (CF_3_SO_2_Na) and 10 mM hydrogen peroxide. The illumination and quenching steps using the FOX photolysis system were performed as previously described.

Desalting and purification were performed using a nonporous graphitized carbon black-solid phase extraction (GCB-SPE) cartridge (Thermo Fisher Scientific, Waltham, MA, USA). The cartridge was preconditioned by sequential washing with 3 mL of 80% acetonitrile containing 0.1% trifluoroacetic acid (TFA) and 3 mL of LC-MS grade water. The sample was then loaded onto the cartridge, washed with 7 mL of LC-MS grade water at a flow rate of approximately 1 mL/min, and eluted with 2 mL of 20% acetonitrile in water. The eluted sample was dried using a lyophilizer and lyophilized sample was reconstituted to yield a final concentration of 25μM in LC-MS grade water, with a portion diluted 1:20 in D_2_O.

For direct infusion analysis, the irradiated and reconstituted samples were subjected to electrospray ionization mass spectrometry (ESI-MS) without chromatographic separation using a Thermo Orbitrap Fusion Tribrid mass spectrometer (Thermo Fisher Scientific, Waltham, MA, USA) equipped with a conductive emitter at a flow rate of 3 μL/min. Data was acquired in positive ion mode using electrospray ionization (H-ESI) with a spray voltage of 400 V, a capillary temperature of 300 °C, and sheath and auxiliary gas flows were set to 1 to ensure stable ion generation and transfer. MS spectra were acquired at a resolution of 60000 and MS^2^ spectra were obtained by collision-induced dissociation (CID) at 30% collision energy. The activation time was set to 10 ms with an activation Q of 0.25. Isotopic ratios at different deuterium incorporation levels were modeled using IsoPro 3.0.

### LC-MS Analysis

All samples were loaded onto a HILIC column (1.0 mm × 150 mm, 1.7 μm, Waters) at a 60 μL/min flow rate coupled to a Dionex Ultimate 3000 nanoLC system Thermo Fisher Scientific, San Jose, CA). Injection volume of the samples was 5µl. Solvent A was 0.1% formic acid in water, and solvent B was 0.1% formic acid in acetonitrile. We eluted the reduced and modified glycans using a gradient consisting of 90% solvent B held constant until 4 minutes, between 4 and 18 minutes, a linear gradient reduces solvent B to 40%, and from 18 to 20 minutes, the B is held constant at 40%, a re-equilibration phase is introduced from 20 to 24 minutes, where the solvent B returns to 90%. The LC column eluant fed into the electrospray ionization source of a Thermo Orbitrap Fusion Tribrid mass spectrometer (Thermo Fisher Scientific, Waltham, MA, USA) using a conductive emitter. All data were acquired in positive ion mode using electrospray ionization (H-ESI) with a spray voltage of 4.0 kV, a capillary temperature of 300 ^°^C, and optimized sheath and auxiliary gas flow for stable ion generation and transfer. MS spectra were acquired at a resolution of 60000, and MS^2^ spectra were acquired in orbitrap mode after collision-induced dissociation (CID) at 30% collision energy, and activation time was set to 10 ms with an activation Q of 0.25. In CID mode, full MS scans were acquired from m/z 200 to 2000, followed by targeted MS^2^ scans corresponding to expected masses of NAG3 and other glycans, along with their modification products.

### Data Analysis

The modified fraction was determined by calculating the ratio of the LC peak areas corresponding to CF_3_ (+67.998) or OH (+15.995) modifications to the total LC peak areas, which included both unmodified and all modified products. Raw MS data files were acquired using Xcalibur software.

We determined the percentage modification using the following equation:

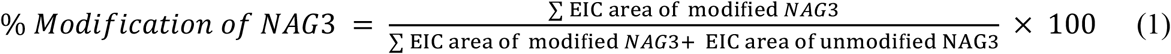

The trisaccharides were identified using extracted ion chromatograms (EIC) of full targeted MS data. We generated fragment masses using GlycoWorkbench and manually assigned MS/MS peaks to theoretical fragments for validation.

## RESULTS & DISCUSSION

### Hydroxyl Radical Carbohydrate Footprinting

To probe the inherent reactivity of carbohydrates, NAG_3_ was used, which consists of three identical n-acetyl-glucosamine monosaccharides with a free-reducing end and a non-reducing end. After oxidation by excimer laser-induced photolysis of hydrogen peroxide, samples were quenched and analyzed. The only oxidation product observed at substantial amounts was the net addition of one oxygen atom (+15.994). No scission products were identified (data not shown). CID MS/MS spectra were obtained. **Figure 1A** shows the annotated MS/MS spectrum of reduced NAG_3_. Reduction of the reducing end allows for differentiation of product ions, and both B- and Y-type ions are observed in the spectrum. Conversely, no masses consistent with reduced (NAG_3_+O) were observed (data not shown). The spectrum of (NAG_3_+O) after attempted reduction **Figure 1B** showed no ions uniquely assignable to the reducing end of the trisaccharide.

**Figure 1.**
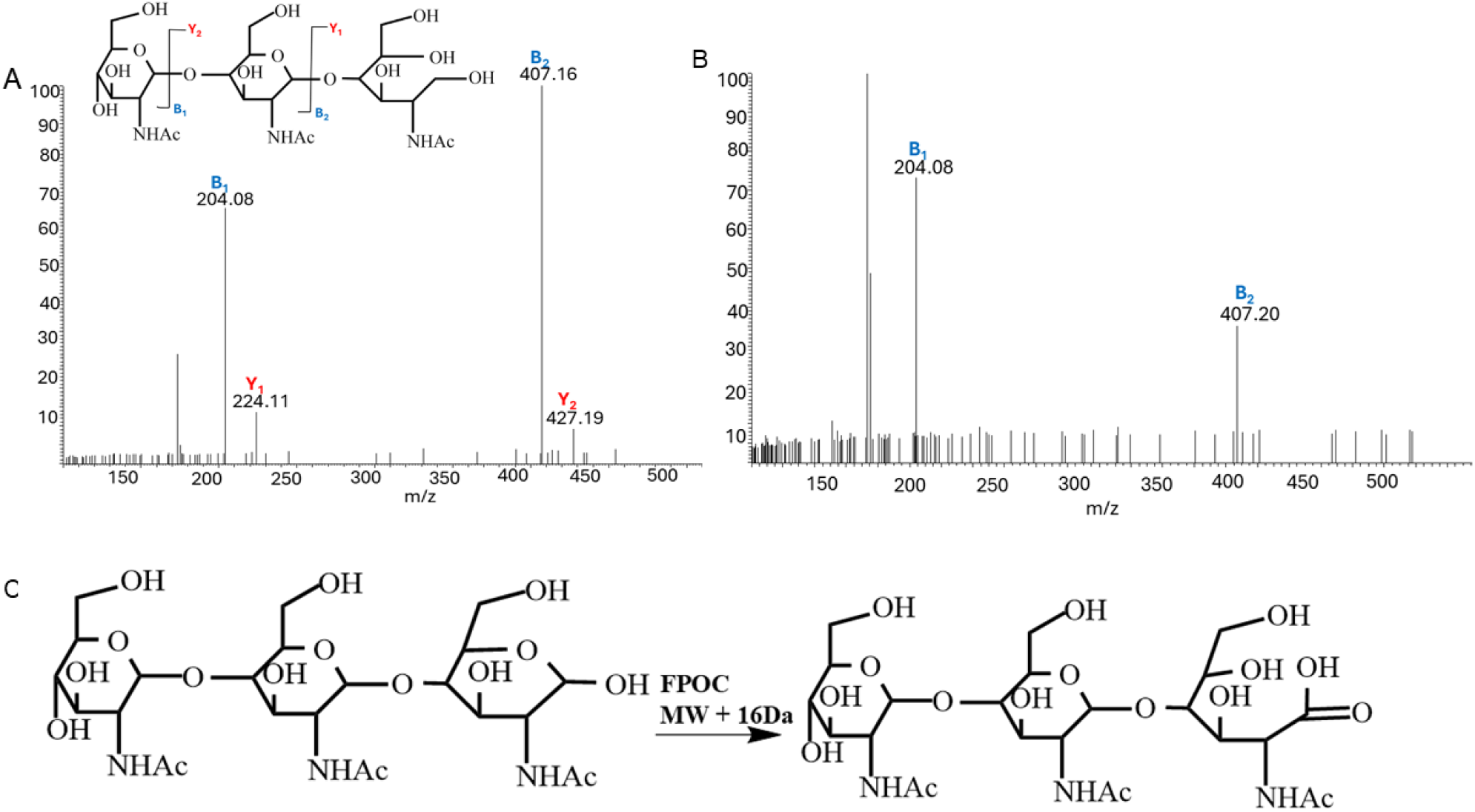
(A) MS/MS spectrum of the reduced NAG_3_ after sodium borohydride reduction. (B) MS/MS spectrum of (NAG_3_+O) after sodium borohydride reduction. No reduced modified NAG_3_ was observed, and no ions containing the reducing end was observed. (C) Proposed scheme for observed hydroxyl radical oxidation product.

In contrast, the B_1_ and B_2_ ions do not show a mass shift consistent with adding an oxygen atom to either of the two monosaccharides at the non-reducing end of NAG_3_. These data together indicate a specific site of hydroxyl radical reactivity at the reducing end. We propose a scheme in **Figure 1C** where the hemiacetal of the reducing end is oxidized by hydroxyl radical to a ring-opened carboxylic acid (which is resistant to sodium borohydride reduction), resulting in the net addition of an oxygen atom.

We then probed the sensitivity of hydroxyl radical reaction rates to changes in solvent accessibility of the reaction site. We used a mixture of five model trisaccharides (NAG_3_, isomaltotriose, 1-kestose, melezitose, and raffinose) and four model proteins (lysozyme, lectin, myoglobin, or ubiquitin) with two non-carbohydrate-binding proteins (ubiquitin and myoglobin) as a control. We used two proteins that specifically bind NAG_3_ and none of the other trisaccharides in the mixture: lysozyme binds the reducing end of NAG_3_[26], while *G. simplicifolia* lectin binds the non-reducing end [27, 28]. We found that non-binding trisaccharides (isomaltotriose, 1-kestose, melezitose, and raffinose) show no significant modification differences in their oxidation event irrespective of the proteins present in the solution, as shown in **Figure 2A**. As shown in **Figure 2B**, the oxidation of NAG_3_ was significantly reduced in the presence of the proteins it binds (lysozyme and lectin) as compared to the samples where non-binding proteins (myoglobin, ubiquitin) are present. Moreover, the reduction in oxidation extent was greater when the protein bound at the reducing end (lysozyme) compared to the non-reducing end (lectin). The results of an ANOVA are presented in **Table S1, Supporting Information**. This protection pattern provides direct evidence of the molecular footprinting capabilities of HRCF, similar to its use in hydroxyl radical protein footprinting [23, 29, 30]. In the absence of protein binding, the modification of NAG_3_ was similar to the modification of the group of isomeric trisaccharides at ∼20%, suggesting that HRCF is a generalizable technique.

**Figure 2:**
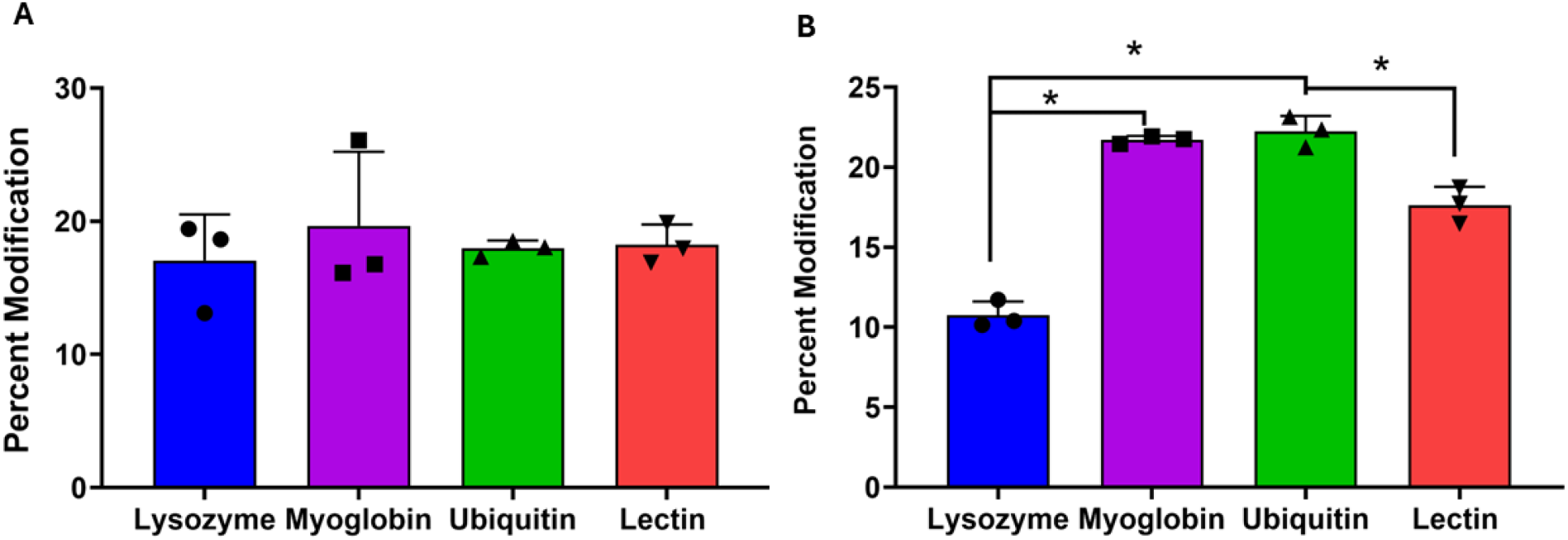
Hydroxyl radical carbohydrate footprinting: An equimolar mixture of NAG_3_ and four isomeric trisaccharides (Isomaltotriose, 1-kestose, raffinose, and melezitose) was oxidized with hydroxyl radicals generated by the laser irradiation of hydrogen peroxide in the presence of different proteins. Brackets with asterisk indicate a statistically significant difference between bracketed pairs by ANOVA with Tukey’s post-hoc analysis (*p* < 0.05). **(A)** The mixture of isomeric trisaccharides that do not bind to any protein show no significant difference in oxidation, irrespective of the protein used. **(B)** NAG_3_ oxidation is significantly reduced in the presence of proteins it binds (lysozyme and lectin) compared to proteins it does not bind (ubiquitin and myoglobin). Error bars represent one standard deviation from a triplicate data.

### Trifluoromethyl Radical Carbohydrate Footprinting

While hydroxyl radicals show utility in footprinting the accessibility of the reducing end of glycans, a more generally reactive radical that can modify glycans at all monosaccharides in the chain would have broader applicability in studying carbohydrate interactions. The Gross lab first introduced the use of sodium triflinate (Langlois reagent) in Fast Photochemical Oxidation of Proteins (FPOP) to generate trifluoromethyl radicals (•CF_3_) for protein footprinting [31]. Subsequently, the Chance lab adapted this strategy to an X-ray radiolysis platform, demonstrating that synchrotron-generated hydroxyl radicals could similarly activate sodium triflinate for trifluoromethyl radical labeling, expanding the toolkit for structural analysis of proteins [21]. This method generates trifluoromethyl radicals from the rapid reaction of sodium triflinate with hydroxyl radicals. Because of the highly different reactivity of trifluoromethyl radicals compared to hydroxyl radicals, we tested the ability for trifluoromethyl radicals to broadly modify our model NAG_3_ glycan, using a commercially available FOX Photolysis System [25] to maximize the ability to replicate the chemistry in other labs.

We first evaluated the effect of different concentrations of hydrogen peroxide on the yield of trifluoromethyl radicals in the presence of 100 mM sodium triflinate and with a FOX system flash voltage of 900V using the mixture of NAG_3_ and isomeric trisaccharides described previously. The only major modification product identified was an increase of 67.987 Da, consistent with hydrogen loss and addition of a trifluoromethyl group. This modification was found on both NAG_3_ and the mixture of trisaccharides, and all glycans modified with CF_3_ were capable of reduction with sodium borohydride (data not shown). As shown in **Figure 3A**, we found maximum modification levels at a peroxide concentration of 40 mM. Interestingly, there was no modification at 10 mM and 20 mM of H_2_O_2_ concentration (data not shown). No difference in modification percentage was found between NAG_3_ and the group of isomeric glycans **Figure 3B**, suggesting trifluoromethyl radicals as is generalizable an RCF label.

**Fig 3.**
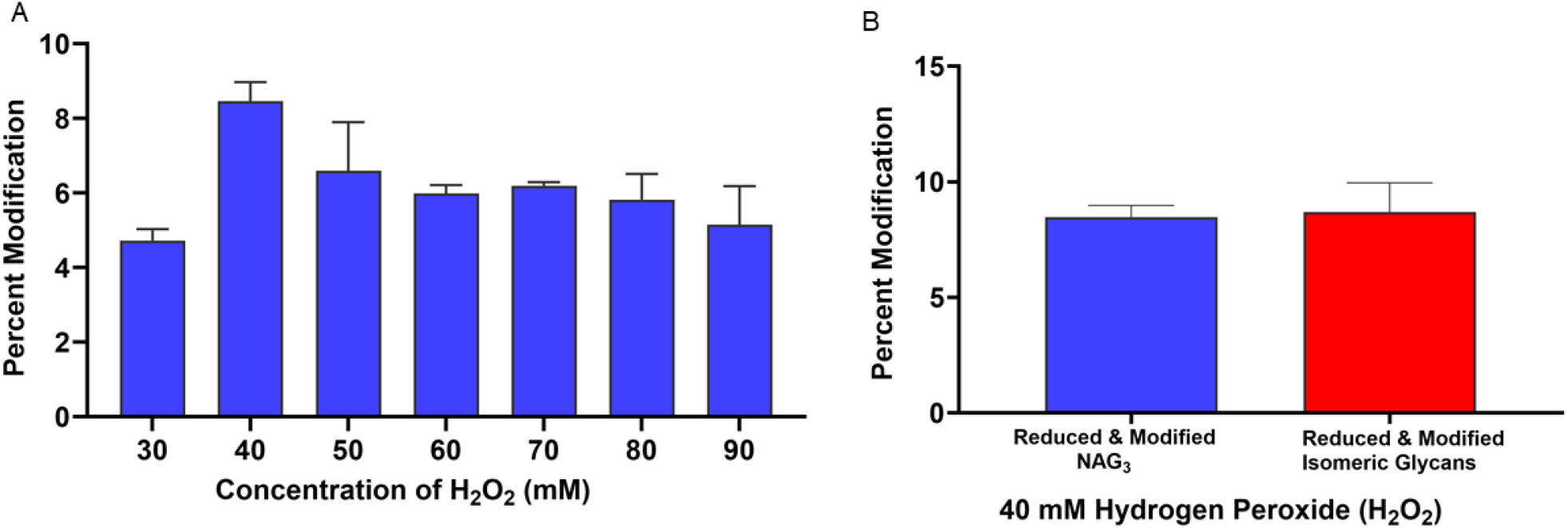
Optimization of H_2_O_2_ concentration for trifluoromethyl RCF. **(A)** Percentage of modified NAG_3_ at different concentrations of H_2_O_2_ in the presence of 100 mM sodium triflinate with a 900V FOX lamp voltage. **(B)** Percentage modification of NAG_3_ and isomeric glycans at 40 mM of hydrogen peroxide.

Next, we tested the sensitivity of trifluoromethyl radical RCF labeling to protein-carbohydrate interactions. The non-binding isomeric trisaccharide mixture shows an apparent reduced modification in the presence of lectin **Figure 4A**, but testing by ANOVA indicates that this apparent reduction is not statistically significant (p = 0.174). **Figure 4B** shows the percent modification of NAG3 with binding proteins (lysozyme, lectin) and non-binding proteins (myoglobin, ubiquitin). The modification of NAG_3_ was reduced in the presence of the two proteins it is known to bind (lysozyme and lectin), indicating structural sensitivity of trifluoromethyl radical RCF. A one-way ANOVA shows that the differences are statistically significant (p = 0.013), with a Tukey’s post-hoc test indicating significant differences between lysozyme and myoglobin (p = 0.012), lysozyme and ubiquitin (p = 0.031), myoglobin and lectin (p = 0.012), and ubiquitin and lectin (p = 0.028). No significant differences were found between lysozyme and lectin (p = 0.999) or myoglobin and ubiquitin (p = 0.871) (**Table S2, Supporting Information)**.

**Figure 4:**
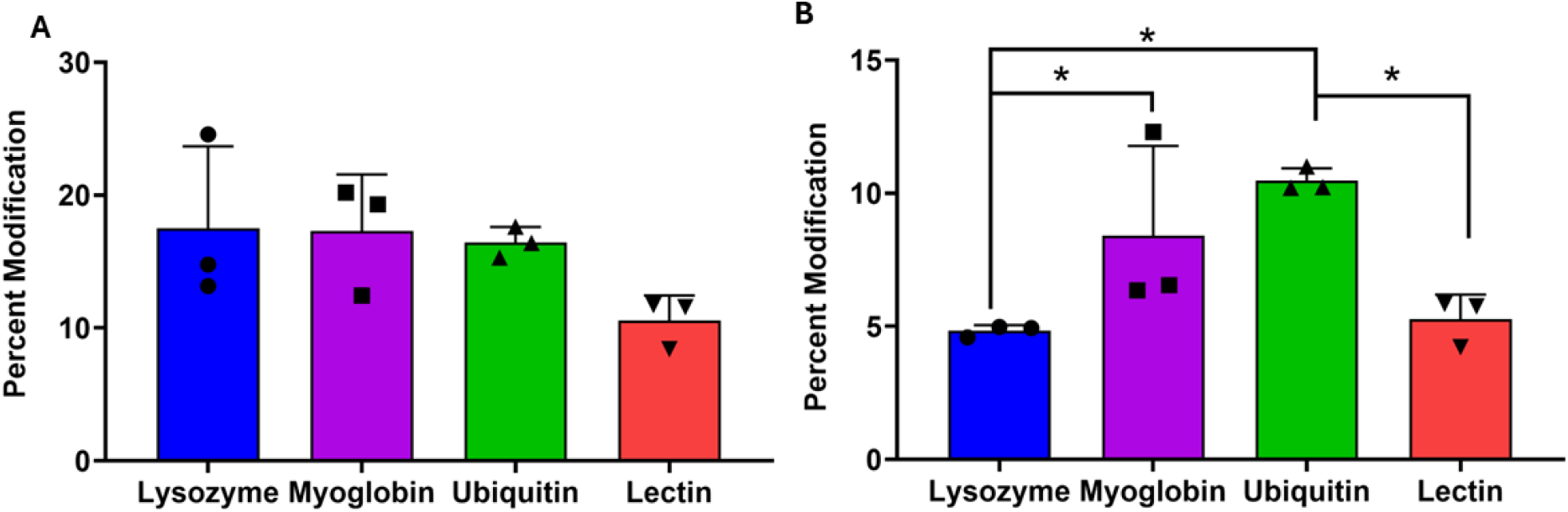
Trifluoromethyl radical carbohydrate footprinting. Mixture of NAG_3_ and isomeric trisaccharides (Isomaltotriose, 1-kestose, raffinose, and melezitose) was oxidized with trifluoromethyl radicals in the presence of a protein. Brackets with asterisk indicate a statistically significant difference between bracketed pairs by ANOVA with Tukey’s post-hoc analysis (*p* < 0.05). **(A)** Isomeric trisaccharides that don’t bind to any protein show no significant difference in modification irrespective of the protein present (p = 0.174). **(B)** NAG_3_ modification is significantly reduced (p<0.05) in the presence of proteins it binds (lysozyme and lectin) as compared to proteins it does not bind (ubiquitin and myoglobin). Error bars represent one standard deviation from a triplicate data set.

MS/MS spectra were obtained to compare unmodified and CF_3_-modified NAG_3_ in both lysozyme and lectin-containing samples. MS/MS spectra were averaged across all partially resolved modification isomers to maximize signal-to-noise. The MS/MS spectrum of unmodified reduced NAG_3_ in the presence of lysozyme is shown in **Figure 5A**. Here, a full b- and y-ion series is present, showing the fragmentation pattern of unmodified NAG_3_. MS/MS of modified reduced NAG_3_ in the presence of lysozyme is displayed in **Figure 5B**. Again, a full b- and y-ion series is present. Each product ion is present in both the modified and unmodified form. Comparison of the relative abundance of the modified to the unmodified version of each product ion (e.g., B_2_:B_2_+CF_3_) clearly shows that the amount of modification increases as the product ion size increases; this is true for both b- and y-ions. These data together indicate that CF_3_ modification is spread across all three monosaccharides of NAG_3_. The MS/MS spectrum of unmodified reduced NAG_3_ in the presence of lectin is shown in **Figure 5C**, with the MS/MS spectrum of modified reduced NAG_3_ in the presence of lectin shown in **Figure 5D**. The MS/MS spectrum of NAG_3_ in the presence of lectin is almost identical to the MS/MS spectrum of NAG_3_ in the presence of lysozyme, suggesting that any differences in the footprint are differences in the quantity of modification at a particular site rather than a complete presence/absence of modification on a monosaccharide. This is consistent with the vast array of data obtained in radical protein footprinting. Unfortunately, while HILIC chromatography gave multiple peaks for reduced modified NAG3, indicating the presence of modification isomers (**Figure 6**), we were unable to identify each modification isomer by MS/MS spectra due to poor MS/MS signal (data not shown).

**Figure 5:**
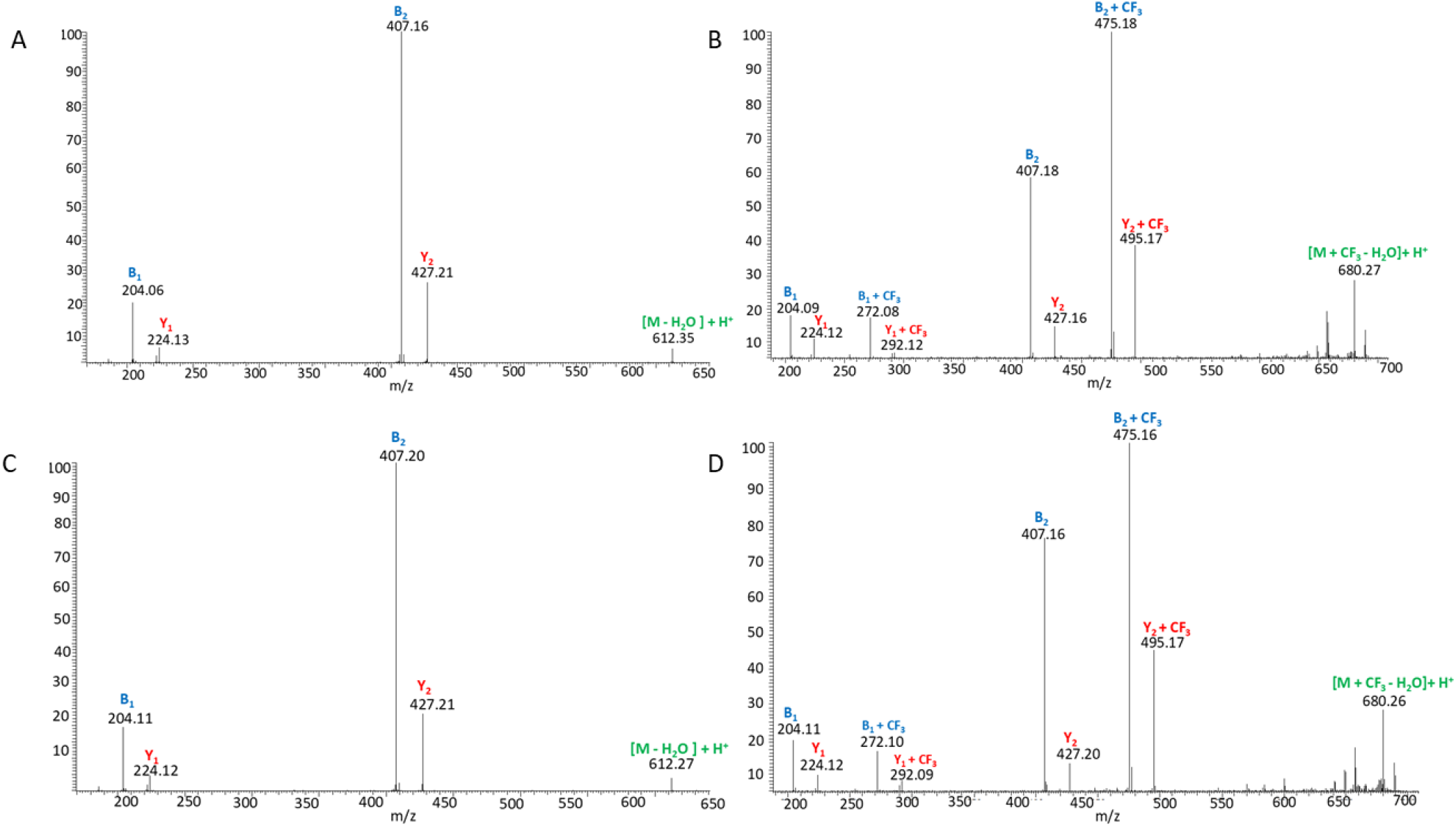
MS/MS spectra of reduced trifluoromethylated NAG_3_. **(A)** MS/MS spectrum of reduced NAG_3_ in the presence of lysozyme. **(B)** MS/MS spectrum of reduced trifluoromethylated NAG_3_ in the presence of lysozyme. **(C)** MS/MS spectrum of reduced NAG_3_ in the presence of lectin. **(D)** MS/MS spectrum of reduced trifluoromethylated NAG_3_ in the presence of lectin.

**Figure 6.**
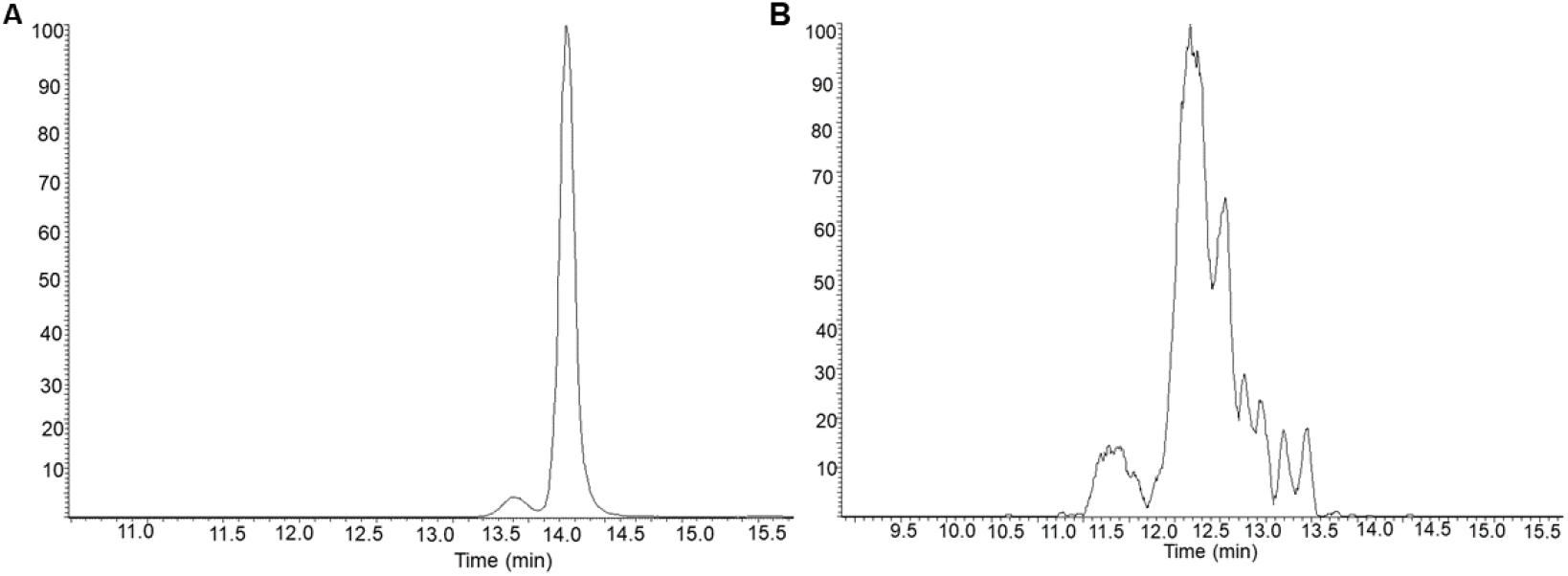
HILIC-MS analysis of trifluoromethylated NAG_3_. **(A)** HILIC-MS extracted ion chromatogram of unmodified reduced NAG_3_. **(B)** HILIC-MS extracted ion chromatogram of trifluoromethylated and reduced NAG_3_.

While our prior data clearly indicates that trifluoromethyl radicals modify glycans on all monosaccharides, it is unclear what molecular mechanism is used to modify the glycan. Two plausible mechanisms exist for modification of NAG_3_ on each monosaccharide; the CF_3_ could replace hydrogen in a C-H bond, or the CF_3_ could replace hydrogen in an O-H and/or N-H amide bond. To investigate the mechanism of trifluoromethyl radical-induced modification on NAG_3_, we used hydrogen-deuterium exchange mass spectrometry (HDX-MS) to differentiate between a non-exchangeable C-H hydrogen replacement from an exchangeable O-H or N-H hydrogen. NAG_3_ contains eleven exchangeable hydrogen atoms, which can be monitored through HDX-MS. Following deuterium exchange, the mass spectrum of unmodified NAG_3_ exhibited the most abundant ion of *m/z* = 656.266, as shown in **Figure 7A**. Isotopic modeling indicated that the mass spectrum shown in **Figure 7A** is most consistent with trifluoromethylated NAG_3_ that retains all 11 exchangeable hydrogens with ∼50% exchange (**Figure S1, Supporting Information)**. The trifluoromethylated ion was observable in the same spectrum (**Figure 7B**). Assuming the same overall deuterium exchange percentage for the unmodified and modified NAG_3_ observed in the same spectrum, the mass spectrum is most consistent with a NAG_3_ ion that retains 11 exchangeable hydrogens (**Figure S2, Supporting Information**). These results indicate that CF_3_ replaces a non-exchangeable hydrogen in a C-H bond rather than an exchangeable O-H or N-H hydrogen to create the final labeled product.

**Figure 7.**
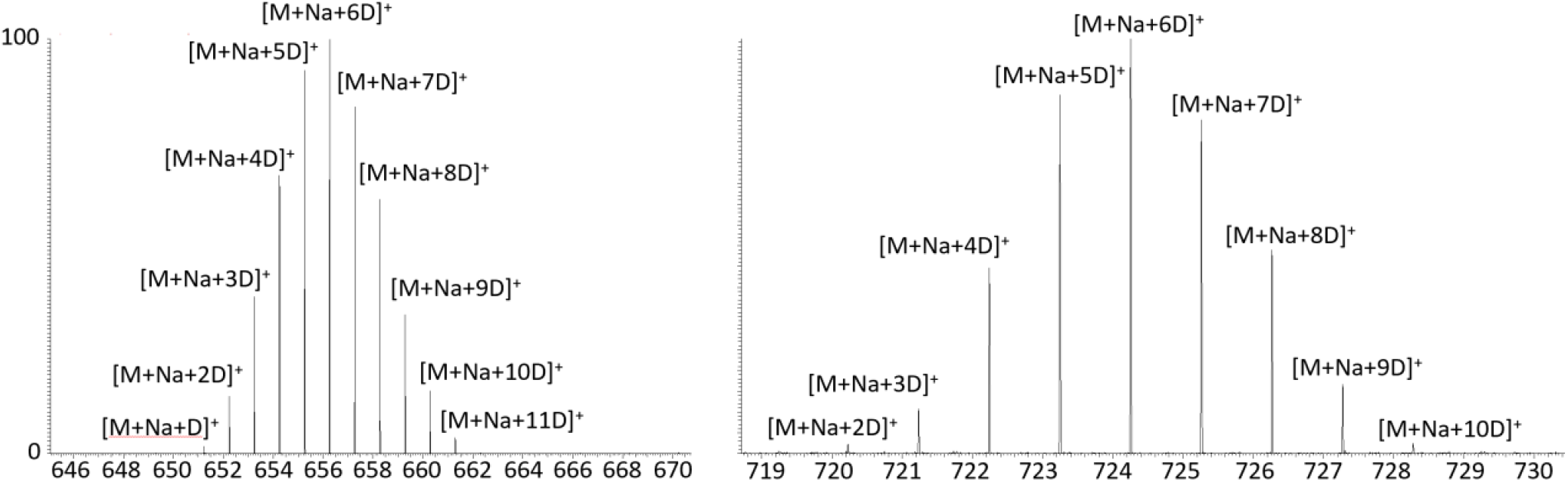
Hydrogen-deuterium exchange shows CF_3_ replaces a C-H hydrogen. **(A)** MS of unmodified NAG_3_ after HDX of ∼50%. **(B)** MS of trifluoromethylated NAG_3_ taken from same spectrum. Spectrum is consistent with a trifluoromethylated NAG3 that retains all 11 exchangeable hydrogens with ∼50% exchange. It is inconsistent with trifluoromethylated NAG3 with only 10 exchangeable hydrogens with ∼50% exchange.

## Conclusion

We demonstrated two separate RCF reagents with their unique purposes: hydroxyl radicals, which specifically label the free reducing end under HRCF conditions, and trifluoromethyl radicals, which modify a variety of glycans across all monosaccharides through substitution at C-H bonds. Both labels exhibited sensitivity to protein-carbohydrate interactions. We were able to measure that the hydroxyl radical is sensitive to the specific portion of the glycan that is recognized by the protein binding partner. We were unable to confirm this with the trifluoromethyl radicals due to difficulties in identifying and quantifying particular positional modification isomers by MS/MS. We are currently working on further advances in properly resolving and identifying these positional isomers of trifluoromethyl radical labeling through improvements in labeled glycan LC separation and improvements in ionization efficiency. These difficulties mirror those encountered in mass spectrometry-based hydroxyl radical protein footprinting [23, 32, 33]; the increased difficulties in ionization of carbohydrates have made RCF challenging, but the hydrophobicity of the CF_3_ group compared to most glycan moieties give us potential options for enrichment and separations to potentially address this issue in future work.

Our observations of the specificity of hydroxyl radical modification are quite different from previously reported results [19], that occurred in hydroxyl radical exposures that were much longer, using lower steady-state concentrations of hydroxyl radicals in a Fenton reaction than the peak concentrations used here in an FPOP-like experiment [34]. While it may be possible to generate a broader HRCF footprint using a slower radical reaction scheme, it is important to remember that these radicals will modify both the carbohydrate analyte and the protein binding partner [35, 36]. Such slow but thorough modification of the protein binding partner can readily change the protein’s conformation [37-39], disrupting protein-carbohydrate interactions and introducing artifacts. Therefore, it is preferable to find a reagent that can broadly modify the carbohydrate analyte under very rapid experimental conditions. Our results indicate that the trifluoromethyl radical is such a reagent. Recent reports from Jain and co-worker [40], introduce the possibility of multiplex trifluoromethyl radical and hydroxyl radical labeling for RCF. Our results in **Figure 3** suggest that modification of the triflinate:hydroxyl radical ratio would adversely affect labeling efficiency, so we chose not to pursue this multiplex approach at this time. However, the approach is intriguing, and particular applications where differentiating labeling at the reducing end from labeling at other regions of the glycan could demand such an approach.

## Acknowledgments

This work was funded by the National Institute of General Medical Sciences (R01GM127267). Research reported in this publication was supported by the Glycoscience Center of Research Excellence, an Institutional Development Award (IDeA) from the National Institute of General Medical Sciences of the National Institutes of Health under award number P20GM130460.

