## Supporting Information for "Hydroxyl and Trifluoromethyl Radical Carbohydrate Footprinting for Probing Protein Binding Components of Oligosaccharides"

**Table S1**. Tukey's post-hoc analysis of ANOVA of •OH RCF in the presence of different proteins

SUMMARY

| *Groups* | *Count* | *Sum* | *Average* | *Variance* |
|---|---|---|---|---|
| Lysozyme | 3 | 32.25 | 10.75 | 0.7213 |
| Myoglobin | 3 | 65.12 | 21.70667 | 0.056633 |
| Ubiquitin | 3 | 66.74 | 22.24667 | 0.920033 |
| Lectin | 3 | 52.91 | 17.63667 | 1.280933 |

ANOVA

| *Source of Variation* | *SS* | *df* | *MS* | *F* | *P-value* | *F crit* |
|---|---|---|---|---|---|---|
| Between Groups | 253.3175 | 3 | 84.43917 | 113.383 | 6.8E-07 | 4.066181 |
| Within Groups | 5.9578 | 8 | 0.744725 | | | |
| Total | 259.2753 | 11 | | | | |

| Group 1 | Group 2 | Mean Difference | p-value | 95% Confidence Interval | Significance |
|---|---|---|---|---|---|
| Lectin | Lysozyme | -6.8867 | <0.0001 | -9.1431 to -4.6302 | Significant |
| Lectin | Myoglobin | 4.07 | 0.0019 | 1.8136 to 6.3264 | Significant |
| Lectin | Ubiquitin | 4.61 | 0.0008 | 2.3536 to 6.8664 | Significant |
| Lysozyme | Myoglobin | 10.9567 | <0.0001 | 8.7002 to 13.2131 | Significant |
| Lysozyme | Ubiquitin | 11.4967 | <0.0001 | 9.2402 to 13.7531 | Significant |
| Myoglobin | Ubiquitin | 0.54 | 0.8673 | -1.7164 to 2.7964 | Not Significant |

**Table S2**. Tukey's post-hoc analysis of ANOVA of •$CF_3$ RCF in the presence of different proteins

**Anova: Single Factor**

| *Groups* | *Count* | *Sum* | *Average* | *Variance* |
|---|---|---|---|---|
| Lysozyme | 3 | 14.50343 | 4.834475 | 0.04621 |
| Myoglobin | 3 | 25.20177 | 8.40059 | 11.42739 |
| Ubiquitin | 3 | 31.44011 | 10.48004 | 0.205678 |
| Lectin | 3 | 15.8214 | 5.2738 | 0.840132 |

| *Source of Variation* | *SS* | *df* | *MS* | *F* | *P-value* | *F crit* |
|---|---|---|---|---|---|---|
| Between Groups | 64.4913 | 3 | 21.4971 | 6.868407 | 0.013259 | 4.066181 |
| Within Groups | 25.03882 | 8 | 3.129852 | | | |
| Total | 89.53012 | 11 | | | | |

### Tukey's HSD Post-Hoc Test Summary:

| Group 1 | Group 2 | Mean Difference | p-Value | 95% Confidence Interval | Significance |
|---|---|---|---|---|---|
| Lysozyme | Myoglobin | -7.11 | 0.012 | -12.52 to - 1.69 | Significant |
| Lysozyme | Ubiquitin | -5.93 | 0.031 | -11.34 to - 0.52 | Significant |
| Lysozyme | Lectin | 0.07 | 0.999 | -5.34 to 5.48 | Not Significant |
| Myoglobin | Ubiquitin | 1.18 | 0.871 | -4.23 to 6.59 | Not Significant |
| Myoglobin | Lectin | 7.18 | 0.012 | 1.77 to 12.59 | Significant |
| Ubiquitin | Lectin | 6 | 0.028 | 0.59 to 11.41 | Significant |

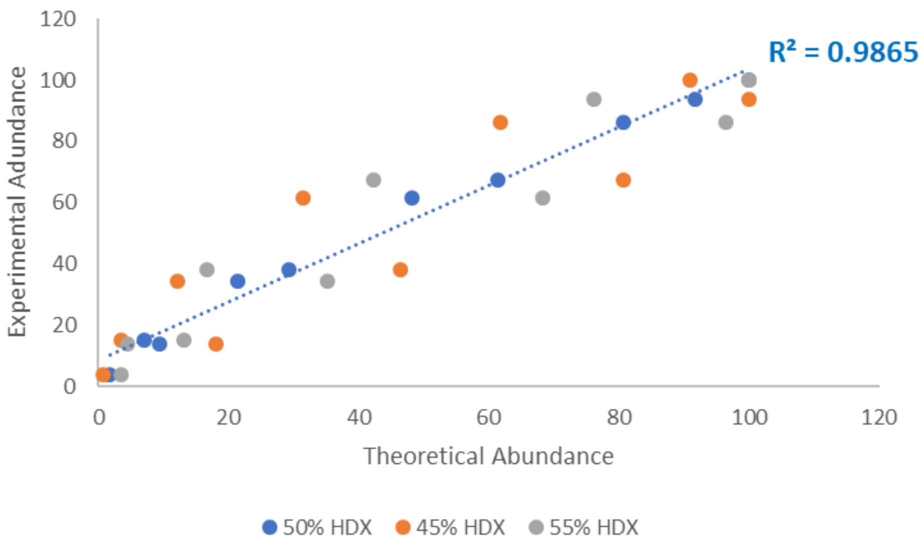

**Figure S1. Correlation of simulated mass spectra with theoretical mass spectra at different deuterium incorporation levels for $NAG_3$.** A deuterium incorporation level of **(blue)** 50% was a substantially better fit ($R^2$ = 0.9865) than **(orange)** 45% deuterium incorporation ($R^2$ = 0.8360) or **(gray)** 55% deuterium incorporation ($R^2$ = 0.8950).

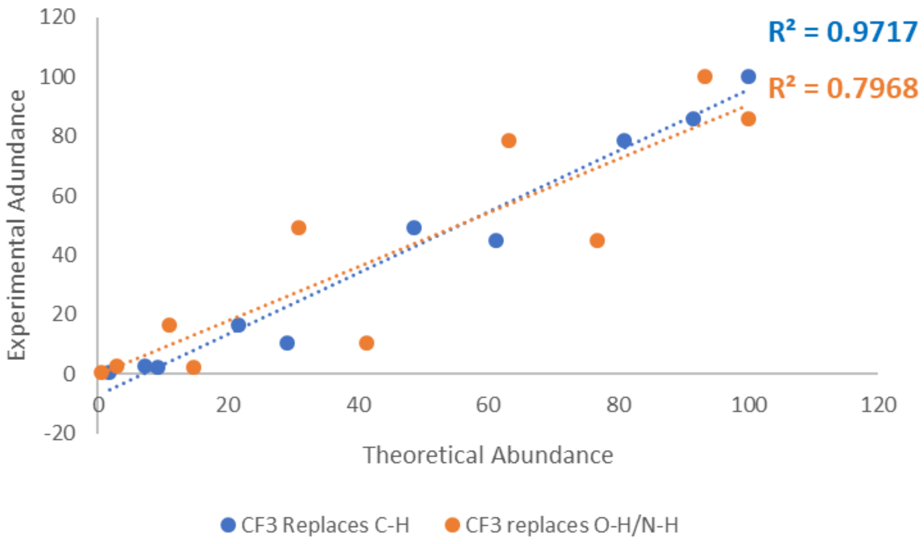

**Figure S2. Correlation of simulated mass spectrum of 50% deuterium exchanged trifluoromethylated $NAG_3$ with experimental results.** The theoretical mass spectrum of trifluoromethylated $NAG_3$ with 50% deuterium content at exchangeable hydrogens matches much better with experimental data when $NAG_3$ has **(blue)** eleven exchangeable hydrogens ($R^2$ = 0.9717) than **(orange)** ten exchangeable hydrogens ($R^2$ = 0.7968).
